# The *K*-mer Antibiotic Resistance Gene Variant Analyzer (KARGVA)

**DOI:** 10.1101/2022.08.12.503773

**Authors:** Simone Marini, Christina Boucher, Noelle Noyes, Mattia Prosperi

**Affiliations:** Department of Epidemiology, University of Florida, Gainesville, FL, USA; Department of Computer and Information Science and Engineering, University of Florida, Gainesville, FL, USA; Department of Veterinary Population Medicine, University of Minnesota, St. Paul, MN, USA

**Keywords:** antibiotic resistance, gene variants, metagenomics, high-throughput sequencing, bioinformatics, statistical learning

## Abstract

Characterization of antibiotic resistance genes (ARGs) from high-throughput sequencing data of metagenomics and cultured bacterial samples is a challenging task, with the need to account for both computational (e.g., string algorithms) and biological (e.g., gene transfers, rearrangements) aspects. Curated ARG databases exist together with assorted ARG classification approaches (e.g., database alignment, machine learning). Besides ARGs that naturally occur in bacterial strains or are acquired through mobile elements, there are chromosomal genes that can render a bacterium resistant to antibiotics through point mutations, i.e., ARG variants (ARGVs). While ARG repositories also collect ARGVs, there are only a few tools that are able to identify ARGVs from metagenomics and high throughput sequencing data, with a number of limitations (e.g., pre-assembly, a posteriori verification of mutations, or specification of species). In this work we present the *k*-mer, i.e., strings of fixed length *k*, ARGV analyzer –KARGVA– an open-source, multi-platform tool that provides: (i) an ad hoc, large ARGV database derived from multiple sources; (ii) input capability for various types of high-throughput sequencing data; (iii) a three-way, hash-based, *k*-mer search setup to process data efficiently, linking *k*-mers to ARGVs, *k*-mers to point mutations, and ARGVs to *k*-mers, respectively; (iv) a statistical filter on sequence classification to reduce type I and II errors. On semi-synthetic data, KARGVA provides very high accuracy even in presence of high sequencing errors or mutations (99.2% and 86.6% accuracy within 1% and 5% base change rates, respectively), and genome rearrangements (98.2% accuracy), with robust performance on ad hoc false positive sets. On data from the worldwide MetaSUB consortium, comprising 3,700+ metagenomics experiments, KARGVA identifies more ARGVs than Resistance Gene Identifier (4.8x) and PointFinder (6.8x), yet all predictions are below the expected false positive estimates. The prevalence of ARGVs is correlated to ARGs but ecological characteristics do not explain well ARGV variance. KARGVA is publicly available at https://github.com/DataIntellSystLab/KARGVA under MIT license.

## I. Introduction

Bacterial antimicrobial resistance is a global threat to human health and to numerous ecosystems, responsible for over 1 million and associated to over 4 million of peoples’ deaths annually (2019 estimate) worldwide [1], disruption of livestock production, and environmental contamination [2]. Resistance in bacteria can manifest naturally, evolve through genetic mutations, or be acquired through gene transfer. While antibiotic susceptibility testing through in vitro cultures is the standard in microbiology research and in clinical/veterinary care settings, high-throughput –targeted and metagenomics– sequencing is becoming a promising alternative, at a relatively low cost and fast turnaround time [3], [4]. The characterization of antibiotic resistance genes (ARGs) from metagenomics as well as cultured bacterial samples through high-throughput sequencing involves development of computational approaches to process large experimental data, up to the terabyte scale, as well as biological annotation of existing and new ARGs into curated database resources [5].

Several public ARG databases are actively maintained, e.g., the comprehensive antibiotic resistance database (CARD) [6], MEGARes [7], the national database of antibiotic resistant organisms (NDARO) [8], ResFinder [9], and the structured antibiotic resistance gene (SARG) database [10], providing access to thousands of gene entries and functional annotations. There is substantial overlap among the databases, however, and it is nontrivial to compare the annotations due to differences in ontology used. For instance, MEGARes uses a tree-based hierarchical structure to represent antimicrobial chemical classes, biological mechanisms, and operon-level gene groups; conversely, CARD uses a reticulate structure. Numerous tools for ARG identification from high-throughput sequencing data are available that rely on one or more of these databases, including methods that are based on: sequence alignment, e.g., AMRPlusPlus [7]; *k*-mers (strings of fixed length *k*), e.g., ResFinder [9], KARGA [11] and AMR-meta [12]; hidden Markov models, e.g., Meta-MARC [13], Resfams [14]; and other machine learning algorithms, e.g., DeepARG [15].

One crucial problem in ARG identification that has gone largely unexplored for metagenomic data is the detection of *ARG variants* (ARGVs) [16]–[18]. ARGVs are constituted by point mutations that allow regular chromosomal genes (e.g., housekeeping genes, topoisomerases) to confer resistance to one or more antimicrobials. It is particularly challenging to detect this type of resistance because both the gene and the mutation(s) must occur. Thus, only a few methods are capable of flagging ARGVs and/or verifying the presence of point mutations. AMRPlusPlus 2.0 flags genes that require variant confirmation, but leaves this additional analysis to the user. CARD’s resistance gene identifier (RGI) identifies ARGVs, but requires partial pre-assembly and protein translation. PointFinder also requires pre-assembly for ARGV identification, and is limited to a number of bacterial strains that have to be specified at runtime. This is a critical issue as ARGVs can encompass a substantial part of ARG databases. For example, 6% of the ARGs in MEGARes are ARGVs.

Here, we present the *k*-mer antibiotic resistance gene variant analyzer (KARGVA). KARGVA is a multi-platform, open-source software specifically designed to confirm the presence of point resistance mutations in chromosomal genes from either metagenomics or whole genome sequencing data. KARGVA merges different ARGV databases and utilizes an efficient three-way, hash-based approach to mutually link point mutations, *k*-mers, and genes. It is equipped with a statistical approach to filter false positives and rank multiple, plausible ARGV candidates. Besides the algorithmic innovation, KARGVA provides advantage over existing tools since it relaxes the need to perform assembly and/or identify species before analysis, and automatically confirms the presence of required mutations during the analysis.

On semi-synthetic data, KARGVA provides high accuracy with sequencing errors or gene mutations and demonstrates robustness with respect to false positive sets. On a large collection of metagenomic data collated by the MetaSUB consortium, KARGVA identifies more ARGVs than AMRPlusPlus, RGI, and PointFinder, and its predictions exhibit a low false positive rate (estimated on a semi-synthetic benchmark set).

## II. Methods

### A. Database Collation

KARGVA integrates ARGV sequences from three different ARG databases: CARD (https://card.mcmaster.ca/), MEGARes (https://megares.meglab.org/), and NDARO (https://www.ncbi.nlm.nih.gov/pathogens/antimicrobial-resistance/). We collect all the protein entries for NDARO. For CARD, we extract ARGVs from the protein variant model, including single resistance variants, multiple resistance variants (with the exclusion of duplications), nonsense mutations, and high confidence *M. Tuberculosis* ARGVs (integrated in CARD from the ReSeqTB database, https://github.com/CPTR-ReSeqTB/UVP). MEGARes ARGVs are compiled through external header and sequence mapping to one of the other sources with available point mutation information. The merged database is made partially non-redundant by collapsing sequences that are identical at the amino acid level. However, multiple mutations are unmodified from the original sources. Thus, if a gene can become resistant through multiple independent point mutations or different combinations of mutations, e.g., the wild-type gene MKRIK mutates into MKRVK or MKRPK to become resistant, all resulting ARGVs appear separately in the database. This is done on purpose to respect the natural occurrence of amino acid variations and their combinations as observed in real-world samples.

Sequence headers and antibiotic resistance ontology metadata, e.g., MEGARes’ class/mechanism/group hierarchy or NDARO’s antibiotic compounds, are pooled together. Since the ontologies of MEGARes, and CARD differ, and NDARO does not use a standardized annotation, KARGVA can report ARGVs with one ontology/annotation term, or more than one if the same ARGV comes from multiple databases.

### B. Classification Approach and Statistical Scoring

The classification of a DNA sequence read as belonging to an ARGV is carried out at the amino acid level, comparing the *k*-mer spectrum of a given protein-translated query sequence with the *k*-mer spectrum of the ARGV database and, separately, of all *k*-mers that include point mutations associated with antibiotic resistance. Since raw high-throughput data is at the nucleotide level, each sequence read is paired to its reverse complement, translated into amino acids using all six reading frames, and then queried against the ARGV database using the algorithm below.

For each query, first it is verified to determine whether its *k*-mer spectrum contains resistance mutations, by comparing it with each ARGV’s subset of *k*-mers that include at least one resistance mutation, out of the complete *k*-mer spectrum of each ARGV. All ARGVs for which at least one positive match of point mutations is found are retained as candidates. Second, the most probable ARGVs are chosen on the basis of the highest relative prevalence of point mutation *k*-mers as well as all *k*-mers that are included in the whole *k*-mer spectrum of an ARGV, but do not necessarily contain point mutations. Since *k*-mers might not be unique to ARGVs, the prevalence scores are weighted by the *k*-mer multiplicity in different genes, e.g., if a *k*-mer is present in three different genes, its weight is one third as compared to a *k*-mer that is found in one gene only. A combined score is the result of the probabilistic sum (P_combined_ = P_1_ + P_2_ – P_1_□P_2_) of the two prevalence measures and decrees the final ARGV ranking. We also use a statistical test that checks how many *k*-mer matches relative to point mutations could be due to chance given a specific sequence length and *k* value. In this way, we are able to filter out false positive classifications at a desired level of confidence. Our procedure calculates the empirical distribution of point mutation *k*-mer occurrences for a sufficiently large (25,000 is the default value) number of random queries, and draws the probability threshold on the basis of the percentile distribution counts, resembling an exact formula previously introduced [19]; in this study, the p-value is set to 0.01. As the ARGV database is only in part non-redundant, there can be cases when more than one ARGV exhibits an optimal score not by chance. To account for these situations, KARGVA reports all scores that are within 5% of the optimal one. Thus, for each sequence read, either no classification to ARGV or classification to one or more ARGVs is provided, with the statistical summaries.

Concomitantly to the per-read classification, KARGVA updates a whole-sample classification, as follows: each time a protein sequence query is assigned to one or more ARGVs, its *k*-mer spectrum is used to update the overall *k*-mer content match for each ARGV in the database. There can be also cases when the *k*-mer content of a read significantly overlaps with that of a region of one or more ARGVs where no required mutations are present, and the read is still used to update the coverage and depth of said ARGV. However, only ARGVs whose resistance mutations are confirmed and pass the statistical filtering are eligible for output, regardless the overall *k*-mer matches. Thus, after processing all reads, KARGVA outputs sample-level coverage and depth summaries for all ARGVs in the database.

### C. Data Structures and Implementation

Three main relational structures are created by indexing: (1) ARGV identifiers linked to *k*-mers (one-to-many); *k*-mers containing point mutations linked to ARGV identifiers (one-to-many); (2) non-mutant *k*-mers linked to ARGV identifiers (one-to-many); (3) ARGV identifiers linked to all their *k*-mers along with frequencies (many-to-one-to-many). KARGVA accepts as input both uncompressed and compressed FASTQ files (recognized automatically through file extension, or as indicated by the user in the command line).

### D. Experimental Setup

Four semi-synthetic datasets are made by simulating high-throughput metagenomic sequencing experiments in different settings, and are used for parameter optimization, performance evaluation and robustness assessment. Specifically, KARGVA is optimized on different values of *k* (between 21 and 45 nucleotides), to determine the best tradeoff between the false positive rate and the false negative rate, with respect to non-ARGV sequences, and ARGV sequences that may carry non-ARGV mutations, gene rearrangements, or sequencing errors.

The first simulated dataset consists of FASTQ files made of: (i) reads drawn from ARGV databases with non-ARGV mutations or sequencing errors up to a 15% rate; (ii) reads from ARGV databases with a two-point transposition and/or transversion, each 50% of the read length; (iii) non-ARG reads, generated uniformly at random. This dataset is used to assess the classification performance on different error rates as well as spurious genes. The second, third, and fourth datasets are simulated using bacterial genomes from the Reference sequence (RefSeq) database at National Center for Biotechnology Information [20], and the Pathosystems Resource Integration Center (PATRIC) web repository [21], where an antibiotic susceptibility test is available. The second read set (semi-synthetic) is generated from 5,000 randomly picked RefSeq bacterial genes (RaBaGe) that did not match any sequence in MEGARes with a BLAST search (e-value=10), putatively susceptible to all antibiotics. The third semi-synthetic dataset is a specific betalactam-susceptible (BeSu) dataset obtained from PATRIC, made by clipping genes of bacterial genomes that (1) are among the top 10% in terms of the numbers of different betalactam antibiotics they were resistant against; and (2) exhibit medium-high similarity to MEGARes genes (BLAST e-value<=0.01, percent identity between 70% and 90%). The fourth semi-synthetic dataset is a specific tetracycline-susceptible (TeSu) dataset collated in the same way as BeSu. In summary, RaBaGe, BeSu, and TeSu are all ‘negative’ datasets (according to the antibiotic susceptibility testing or (non)match with MEGARes) on which none of the classifiers should find antibiotic resistance genes, unless the genes were present but not expressed. RaBaGe covers the spectrum of all genes, while BeSu focus on genes that are very similar to ARGVs but present different mutations. With all datasets we assess and optimize the false positive rate of KARGVA by varying *k* and gene coverage threshold. The semi-synthetic datasets are simulated using InSilicoSeq software with presets for Illumina [22].

We finally evaluate KARGVA on real experimental data, using metagenomic experiments from samples collected by the MetaSUB consortium [23]. MetaSUB is an international project that collects surface samples from public transportation systems, and collates metagenomic sequencing of urban microbiomes with the goal *“to build a molecular profile of cities around the globe to improve their design, functionality, and impact on health*.*”* We replicate in part the work presented by Danko *et al*. [24], who applied AMRPlusPlus to detect both ARGs and ARGVs, and provided an abundance, density and correlation analysis among sampled cities’ metagenomes in relation to their distance and distribution of resistance genes. To extract the species present in each sample, we utilize Kraken2 [25] and its standard database version as of 2019-09-04. In addition to correlation and density maps for sample characteristics by ARGs and ARGVs, we fit a random forest (RF) to evaluate how much variance in ARGVs is explained by the sample attributes – the city of origin, surface material (manually curated, grouping materials that are not present in at least 5 cities and in a total of 25 samples into an ‘other’ category), type of sample (air, environmental, unknown), number of total reads per sample, and bacterial species, classified with Kraken2. We use R (https://www.r-project.org/) with packages caret and ranger, optimizing the number of trees and number of splits at each tree node via grid search. RF variable importance is evaluated via permutation. Prediction performance is assessed through 10-fold cross validation, measuring the root mean squared error (RMSE) and mean absolute error (MAE).

We further compare KARGVA with AMRPlusPlus 2.0, RGI 5.1.1 and PointFinder 4.2. AMRPlusPlus is an alignment-based method that identifies ARGs as well as candidate ARGVs (that require confirmation of mutations) from high-throughput read data using MEGARes. RGI is also alignment-based, but it uses protein-translated queries and CARD. PointFinder is a hybrid method that uses alignment and *k*-mers. Differently from KARGVA and AMRPlusPlus, RGI and PointFinder require whole genomes or genes. Thus, in order to evaluate RGI and PointFinder, the raw read files are assembled using metaSPAdes 3.15.3 (recommended parameters), after quality control, filtering and adapter trimming preprocessing [26]. Furthermore, as PointFinder requires to specify an input species, we run it for all the supported species, merging its output and removing duplicates. For comparing ARGV predictions in this work –since ARGV classification tools may use different ontology– we manually review all the antibiotic resistance ontology/prediction annotations of KARGVA, RGI, and PointFinder, using MEGARes classes as a common reference. All terminology for which we are able to find a correspondence is retained (see Supplementary Material).

All tests are run on University of Florida’s HiPerGator computing cluster (https://www.rc.ufl.edu/) on nodes with 4 Intel Xeon CPUs at 2.00 Ghz with 16 GB RAM.

## III. Results

The KARGVA ARGV database integrates 1,159 single-point, and 95 multiple-point ARGVs from CARD (Jan 2021 release); and 654 single-point ARGVs from NDARO 3.10, respectively. In MEGARes, ARGVs are not provided with information on the variant location, but we are able to align 1,173 MEGARes’ ARGVs with CARD, and 147 with NDARO. After merging identical sequences, the final KARGVA database includes 1,781 ARGVs. Out of the total, 95.5% of sequences contain just one point mutation conferring antibiotic resistance, 3.3% contain two, 1.1% contain three or four, and 2.5% contain stop codons. KARGVA reports ARGV annotations in accordance with all original database ontologies; in the case of a sequence that is present in more than one database, all individual annotations are provided together.

The first semi-synthetic dataset is used to optimize the *k* value and to assess the single best match performance across different sequencing error rates (or more in general as any nucleotide change from the original ARGV sequence) and gene rearrangements. Each simulation comprises 25,000 reads of length of 151 bases. **Figure 1** (left) displays the accuracy curves (with 95% confidence intervals drawn across simulations) stratified by *k* value. Larger *k* values are more accurate at lower error rates, and vice-versa. The best tradeoff is given by *k*=9 amino acids (i.e., 27 nucleotides), yielding accuracies well over 80% for error or base change rates up to 2.5%. **Figure 1** (right) illustrates the distribution of scores for the single best match across all experimental configurations, stratified by the classification correctness. The distributions clearly indicate that the proposed score has very high discriminative ability, yielding a median (IQR) value of 0.83 (0.59-0.99) for the correctly identified genes vs. 0.41 (0.21-0.60) for the wrongly/non-identified entries. As explained, more than one ARGV can have optimal score, so we can evaluate all best scoring ARGVs. **Figure 1** (middle) shows the performance of our method using the best *k* and the top scoring matches. The accuracy is 99.2% for error rates up to 1%, 93.5% at 2.5%, 79.8% at 5%, 50.3% at 10%, and 26.9% at 15%. For 2-point gene rearrangements, the overall accuracy is 98.2%.

**Figure 1.**
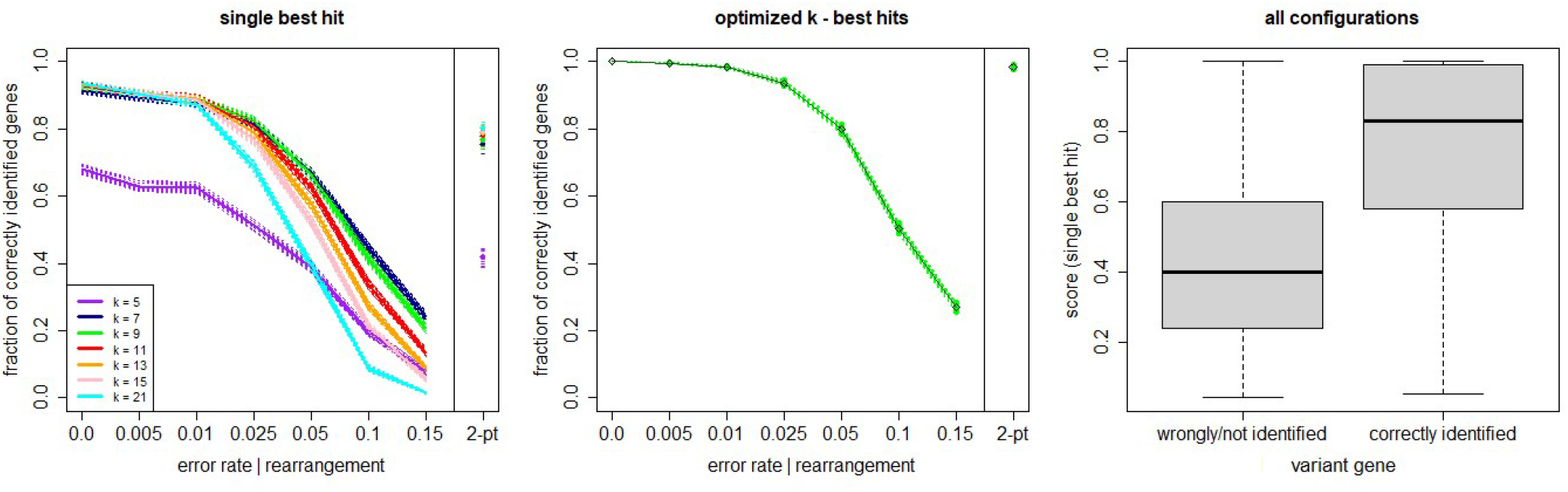
KARGVA performance on semi-synthetic data (varying error / nucleotide change rates and gene rearrangements) in identifying point mutations in chromosomal genes of bacteria conferring antibiotic resistance. Left panel shows performance for a single best match, stratified by parameter *k* value; Middle panel shows performance after *k* optimization, using all optimal best scoring matches; right panel shows box-and-whisker plot distribution of algorithm’s scores on all test configurations, stratified in accordance with ground truth. Shaded areas and whiskers represent 95% confidence intervals.

The other three semi-synthetic datasets help us determine the robustness of KARGVA with respect to false positive rate on more realistic bacterial metagenomics data. For RaBaGe, we select over 5,000 genes with length >500 nucleotides that meet the antibiotic susceptibility criteria, and generate 500,000 reads, for BeSu we obtain 4.2 million reads, while for TeSu 355,170 reads. We benchmark KARGVA over multiple parameter combinations, varying the minimal required gene fraction coverage between 0.00 and 0.95, and considering *k* equal to 7, 9, 15 (i.e., 21, 27, 45 nucleotides). For each parameter configuration and dataset, we calculate the false positive score (FPS), measured as # of detected ARGVs / # of reads. As shown in **Figure 2**, even at low coverage thresholds and small *k* values, the FPS is low. With any coverage fraction and *k*=7, the FPS is 3.7×10^−4^ in RaBaGe, 1.5×10^−4^ in BeSu, and 1.7×10^−4^ in TeSu. With the most conservative coverage of 0.95 and *k*=15 (i.e., long conserved stretches), FPSs is 2×10^−6^, 3.8×10^−6^, and 1.1×10^−5^ for RaBaGe, BeSu, and TeSu, respectively. We then set the KARGVA to default configuration with coverage of 0.8 and *k*=9, which yields FPS rates all below 5 in 100,000, specifically 4×10^−6^, 4.5×10^−5^, and 4.2×10^−5^ for RaBaGe, BeSu, and TeSu. We expect real metagenomics data to be in-between the ‘easy’ (RaBaGe) and ‘hard’ (BeSu, TeSu) negative datasets, and therefore we estimate the expected KARGVA FPS on metagenomics data to be in the [10^−5^, 10^−6^] range when used with default settings. We also test RGI and PointFinder on these three negative datasets. RGI shows a worse FPS, yielding at least twice false positives than KARGVA. Over RaBaGe, BeSu, and TeSu, RGI FPSs are 6×10^−6^, 9×10^−5^, and 1.2×10^−4^, while PointFinder shows the best FPSs with 4.8×10^−7^, 3.3×10^−6^, and less than 2.8×10^−6^, i.e., no findings over TeSu. Note that both RGI and PointFinder are run on contigs assembled by metaSPAdes, and not on the original read sets.

**Figure 2.**
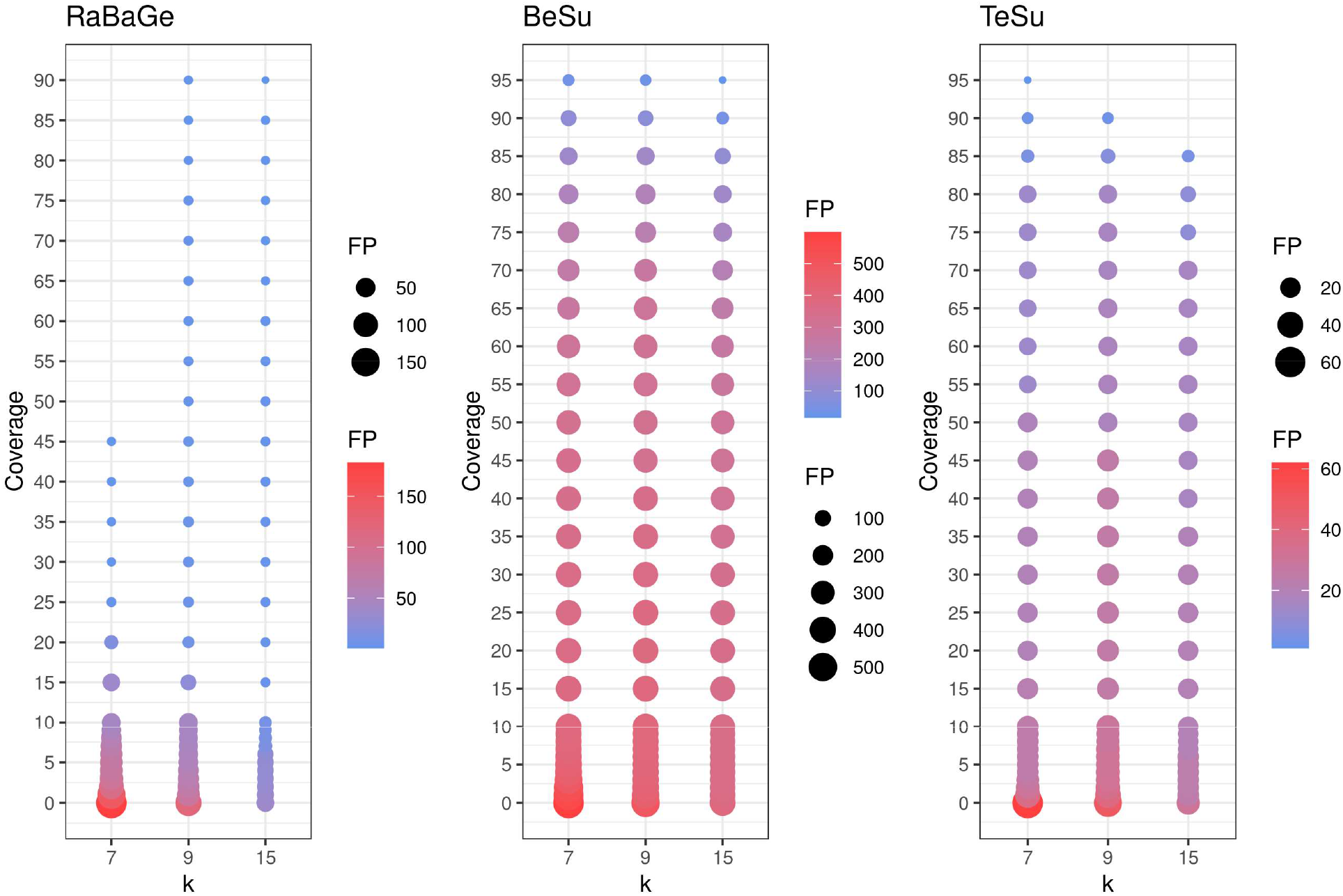
Assessment of KARGVA’s false positive rate on semi-synthetic data by varying *k*-mer length and gene coverage threshold. The X axis represents the *k* value (in amino acids, e.g., *k*=9 means 27 nucleotides), the Y axis represents the gene coverage portion (%), and the bubble size represent the false positive counts. RaBaGe, BeSu, and TeSu are synthetic datasets assembled by random bacterial genes (RaBaGe), and PATRIC genome fragments exhibit ingmedium-high similarity to MEGARes betalactamase (BeSu) or tetracycline (TeSu) genes.

Next, we run KARGVA, AMRPlusPlus, metaSPAdes+RGI and metaSPAdes+PointFinder on the metagenomics global data (surface samples from public transportation) from the MetaSUB project. The MetaSUB FASTQ files available for public download are filtered for human DNA. Of 4,305 paired short read files, 3,758 come with a matched metadata record and belong to a city with at least 25 samples, and can thus be processed (i.e., provide an output) by all the algorithms we used. **Table 1** shows sample/isolate characteristics for the MetaSUB files selected and analyzed, considering the top-10 cities in terms of total number of samples, with summaries of the top-5 most frequent species as classified by Kraken2, and the mean, median (interquartile) number of ARGs detected by AMRPlusPlus and of ARGVs detected by KARGVA. **Figure 3** shows the relationships between species abundance and city, considering the top-5 species. Out of 5,053 unique species detected, 22 make the top-5. *Cutibacterium acnes* is the most abundant in terms of average per-sample reads in eight of the top-10 cities. Of note, a considerable fraction of the reads (28%-55%) cannot be assigned to a species using the standard Kraken2 database.

**Table 1.** Summary of MetaSUB study data stratified by city (top-10 by number of isolates, others aggregated). We report the 5 most frequent bacterial species from each location (based on per sample prevalence), along with the median percentage of unclassified samples, as found by Kraken 2.0. Antibiotic resistance genes (ARGs) are identified using AMRPlusPlus 2.0, while ARGVs are identified using KARGVA. IQR: interquartile range.

| City | No. of samples | Top-5 species | Mean; Median (IQR) no. of ARGs per sample | Mean; Median (IQR) no. of ARGVs per sample | Median (IQR) % of unaligned reads per sample |
| --- | --- | --- | --- | --- | --- |
| Hong Kong | 770 | *Cutibacterium acnes*; *Bradyrhizobium* sp. BTAi1; *Micrococcus luteus*; *Cupriavidus metallidurans*; *Janibacter indicus* | 21.28; 17 (9, 28) | 3.9; 2 (2, 3) | 33.02 (26.59, 42.87) |
| London | 635 | *Cutibacterium acnes*; *Staphylococcus simulans*; *Micrococcus luteus*; *Kocuria rosea*; *Staphylococcus aureus* | 18.57; 4 (1, 14) | 23.85; 2 (2, 4) | 42.18 (32.85, 48.41) |
| New York City | 493 | *Cutibacterium acnes*; *Pseudomonas stutzeri*; *Micrococcus luteus*; *Stenotrophomonas* sp. LM091; *Kocuria indica* | 36.92; 20 (9, 41) | 25.01; 4 (2, 23) | 28.14 (18.42, 43.07) |
| Ilorin | 264 | *Pseudomonas stutzeri*; *Cutibacterium acnes*; *Pseudomonas balearica*; *Bradyrhizobium* sp. BTAi1; *Acinetobacter* sp. ACNIH1 | 57.9; 31.5 (9.75, 75.75) | 36.5; 13 (2, 56.5) | 43.38 (25.79, 55.16) |
| Singapore | 179 | *Cutibacterium acnes*; *Bradyrhizobium* sp. BTAi1; *Micrococcus luteus*; *Bradyrhizobium* sp. SK17; *Kocuria rosea* | 18.58; 9 (4, 17.5) | 12.75; 2 (2, 4) | 43.19 (32.68, 54.12) |
| Tokyo | 148 | *Cutibacterium acnes*; *Bradyrhizobium* sp. BTAi1; *Cupriavidus metallidurans*; *Bradyrhizobium* sp. SK17; *Moraxella osloensis* | 40.29; 13.5 (6, 32.5) | 26.23; 2 (2, 8) | 32.53 (22.33, 55.41) |
| Barcelona | 116 | *Acinetobacter junii*; *Staphylococcus aureus*; *Enterococcus faecium*; *Cutibacterium acnes*; *Salmonella enterica* | 71.53; 10 (2.75, 24.5) | 91.7; 4 (2, 15) | 36.49 (18.89, 59.49) |
| Stockholm | 110 | *Cutibacterium acnes*; *Bradyrhizobium* sp. BTAi1; *Micrococcus luteus*; *Pseudomonas stolaasi*; *Kocuria rosea* | 5.18; 2 (0, 6) | 2.07; 2 (2, 2) | 54.55 (48.58, 58.99) |
| Porto | 107 | *Cutibacterium acnes*; *Staphylococcus simulans*; *Staphylococcus epidermis*; *Cupriavidus oxalaticus*; *Staphylococcus aureus* | 5.8; 2 (0, 6) | 23.52; 2 (2, 2) | 53.11 (45.77, 61.76) |
| Fairbanks | 95 | *Cutibacterium acnes*; *Staphylococcus epidermidis*; *Micrococcus luteus*; *Moraxella Osloensis*; *Staphylococcus haemoliticus* | 26.44; 13 (4, 35.5) | 7.59; 2.5 (2, 15) | 43.55 (25.61, 55.05) |
| Other (aggregated) | 841 | *Pseudomonas stutzeri*; *Cutibacterium iumacnes*; *Bradyrhizobium* sp. BTAi1; *Acinetobacter woffii*; *Moraxella osloensis* | 32.58; 11 (4, 25) | 20.02; 2 (2, 5) | 45.44 (32.16, 53.57) |

**Figure 3.**
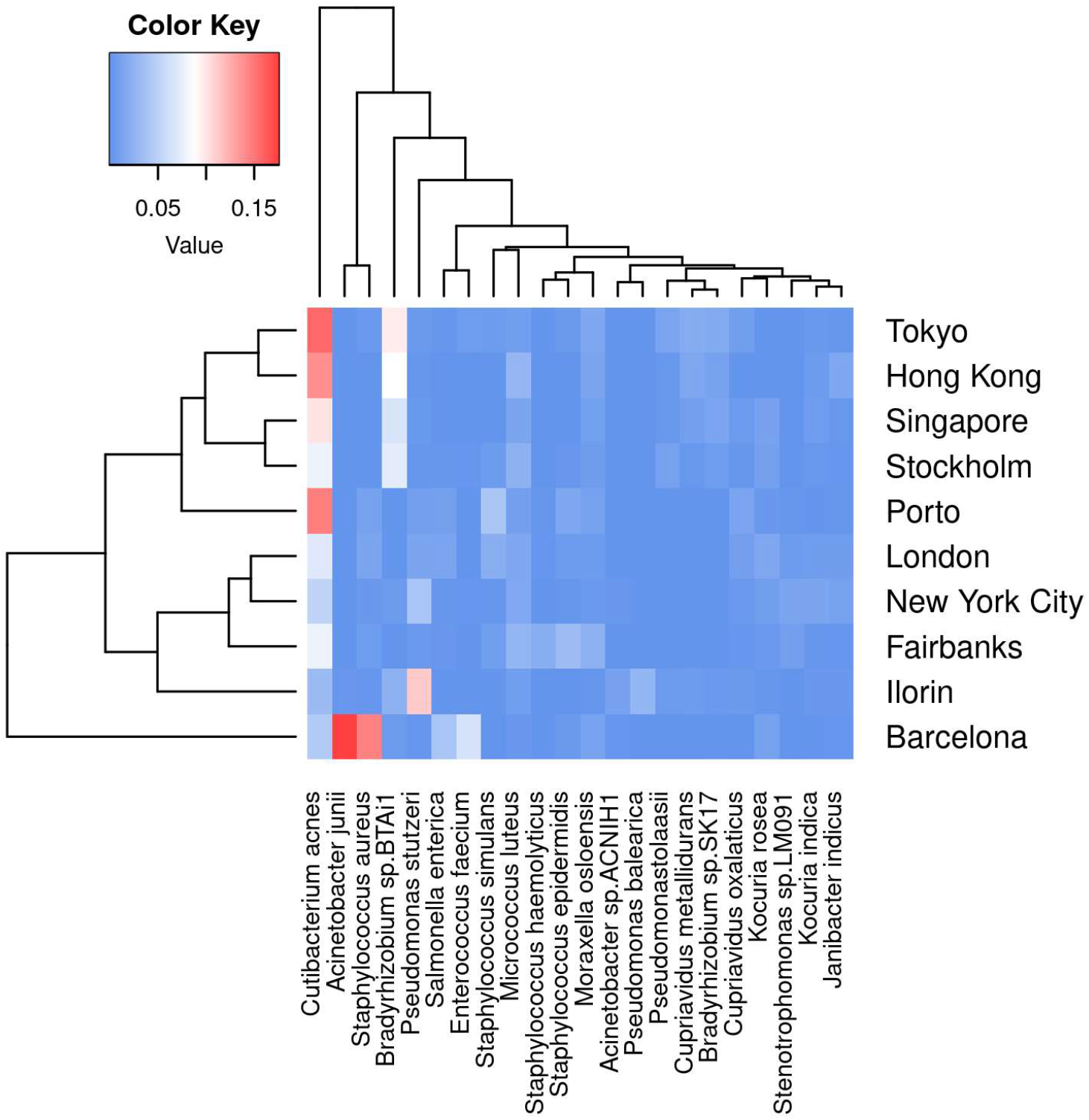
Average per-sample frequency of bacterial reads, considering the top 10 cities per number of sample, and the top-5 species per city. The color key represents the prevalence in [0,1] scale of each species.

In order to assess the reliability of ARGV findings, we compare in detail the ARGV detection by KARGVA with respect to that of AMRPlusPlus, RGI, and PointFinder. In **Figure 4**, we calculate ARGV counts for all algorithms overall (i.e., total number of ARGVs, independently from the class) and per-class. Through the ontology/annotation mapping described in the methods, we identify eleven MEGARes classes that can be predicted by all three algorithms: amingoglycosides; betalactam; fluoroquinolones; fusidic acid; lipopeptides; macrolide, lincosamide and streptogramin (MLS); mupirocin; oxazolidnone; rifampin; sulfonamides; and tetracyclines. For this reason, the per-class comparison must be limited to the classes all algorithms can predict. Of note, while by design KARGVA has a single MEGARes class assigned to each prediction, RGI and PointFinder might have multiple, i.e., multiple output terms can match the same MEGARes class in a sample. We therefore allow RGI and PointFinder to count multiple times if the annotation terms of their predictions match with more than one class (see Supplementary Material). Overall, KARGVA finds 43,846 ARGVs, ∼4.8 times more than PointFinder (9,185) and ∼6.8 more than RGI (6,472). KARGVA retrieves the highest number of ARGVs in 8 out of 11 considered classes, the exceptions being aminoglycosides (highest: PointFinder), MLS (highest: RGI), and Oxazolidinone (highest: PointFinder). Although it is not possible to directly transpose the FPS from the synthetic datasets to the MetaSUB results, we expect RGI to find more false positives then the other two algorithms, and PointFinder to be the most conservative. For a reference, AMRPlusPlus yields over 100,000 ARGVs that need SNP confirmation; KARGVA, RGI, and PointFinder are all well below this value.

**Figure 4.**
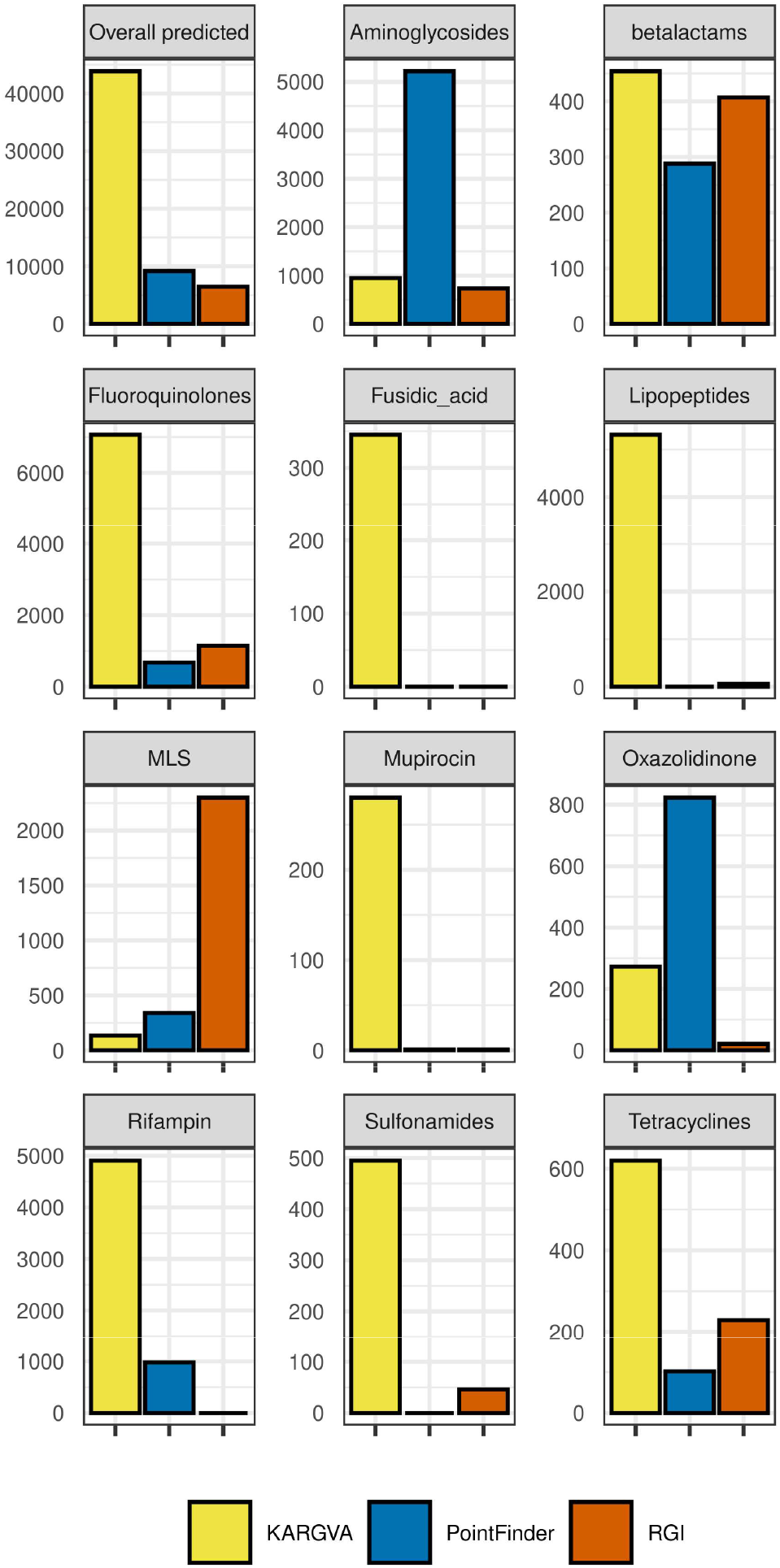
ARGVs detected in the MetaSUB metagenomics datasets (*n*=3,758 with available metadata) by KARGVA, RGI, PointFinder, both overall and per class.

We then analyze how KARGVA’s ARGVs relate to the ecological characteristics of the samples and the ARG distributions. Using the top-10 city set, the per-sample median and mean number of retrieved ARGs by AMRPlusPlus (mean range: 2-31.5; median range: 5.18-71.53) are comparable to the retrieved ARGVs by KARGVA (mean range: 2-13; median range: 3.9-91.7), and they are strongly correlated (Spearman’s correlation for mean and median, respectively: 0.63; 0.89). Note that we cannot achieve a perfect replication of the original ARG analysis presented by Danko *et al*. [24], since the original MetaSUB analysis used MEGARes 1.0.1 and Bowtie 2.3.0. Instead, we apply filtering criteria on the cities based on sample size, and use the most up-to-date AMRPlusPlus 2.0 pipeline, which employs MEGARes 2.0 and BWA, along with specific preprocessing (Trimmomatic) and post-processing (Bedtools, SNPfinder), finalizing with the ResistomeAnalyzer. Nonetheless, there is consistency in the overall output, with more positive identifications expected, since MEGARes 2.0 contains more genes than the prior release.

By stratifying the distribution of ARG and ARGV findings per city and antibiotic class, we check if there are relevant correlations between city and class or between classes. **Figure 5** shows: the per-city ARG and ARGV (panels B and E) distributions, the per-city ARG and ARGV (panels A and D) class profiles –defined as the fraction of the per-class counts over the total counts of a city– and the per-class correlation (Spearman) based on the ten city profiles. Large fractions of ARG counts per-city come from MLS, betalactam, and aminoglycoside classes, while the highest fractions of ARGV counts come from fluoroquinolones. The class-to-class correlation structures are different between ARGs and ARGVs. For instance, fluoroquinolones and lipopeptides, as well as MLS and aminoglycosides ARGs are found often together, while the correlation is low in ARGVs, where fluoroquinolones and aminoglycosides tend to cluster apart from the others. Of note, there are differences among the ARG/ARGV classes reported. Some resistance classes, such as Oxazolidinone, are not present as ARGs, since MEGARes and KARGVA annotations have only a partial overlap.

**Figure 5.**
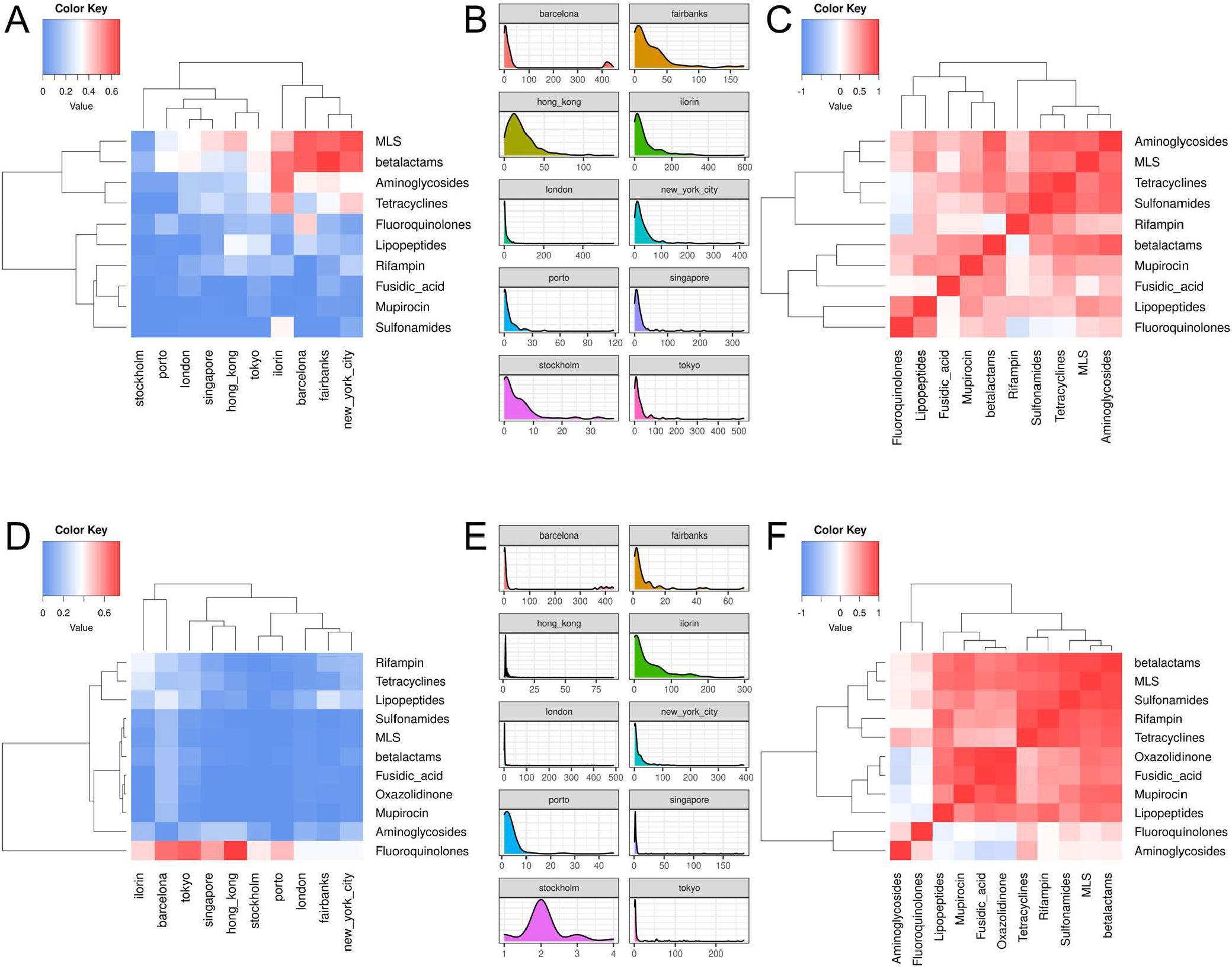
Correlation heatmaps (fraction of counts per antibiotic resistance class over the city total) and density plots of antibiotic resistance genes (ARGs, Panels A-B-C) and ARG variants (ARGVs, Panels D-E-F), among the top-10 cities and resistance types using the MetaSUB annotated samples (n=2,917).

After correlation and density analysis, we fit the RF model to predict the number of ARGVs per samples given the city of origin (24 cities), surface material, type of sample, total reads, and bacterial species (5,054 species by Kraken2). Using 250 trees and varying the number of attributes considered at each split, performance varies from 34.6±1.41 (RMSE) and 14.2±1.41 (MAE) with 2 splits per node, to 12±2.3 (RMSE) to 4.3±0.38 (MAE) with 5,089 splits per node, i.e., considering all the variables at each split like in a regular decision tree bagging algorithm. Both RMSE and MAE can be considered very high compared to the median numbers of ARGVs found in each sample, i.e., 2 to 13 across all cities. In terms of variable importance, the number of ARGs holds the highest predictive value, which matches the high univariate correlation of this feature with the number of ARGVs, followed by bacterial species (top-3 are: *Rahnella spp*. Y9602, *Chlamydia spp*. 2742.308, and *Chlamydia gallinacea*); the first non-species variable is the number of reads, ranked 24^th^.

In regards to data processing speed, we compared KARGVA and AMRPlusPlus on a benchmark obtained by clipping a MetaSUB sample (haib17CEM4890_H2NYMCCXY_SL254773) to 12.5, 6.25, and 3.125 million read pairs (approximatively 8, 4, and 2 GB). KARGVA processes on average 1GB of FASTQ data in 00:02:30 hh:mm:ss with processing times of 00:04:31, 00:09:37, and 00:17:52 for, respectively, 2, 4 and 8 GBs inputs. Importantly, KARGVA memory usage is extremely contained, requiring less than 512MB RAM regardless of the FASTQ size (with 150-300 bp read lengths). We noticed the processing time linearly increasing with file size and the difference between processing compressed vs. uncompressed files is minimal. On the other hand, AMRPlusPlus is about 6 times slower, with processing times of 00:31:16, 01:01:31, and 01:59:26 for, respectively, 2, 4 and 8 GB inputs. Of note, we could not properly compare processing times against RGI and PointFinder as they do not process FASTQ inputs directly, and require pre-assembly, which is computationally intensive. For instance, the whole assembly of the MetaSUB data requires about a month of wall time (including job scheduling operations) on the HiPerGator cluster, with over 1,000 normalized computing units.

## IV. Discussion

KARGVA is a fast, multi-platform software for detection of ARGVs from metagenomic high-throughput sequencing experiments. Its ARGV database integrates multiple sources and is the largest available to date, with full linkage to the original sources and their respective ontologies for annotation of resistance class and mechanisms. Our method confirms the detection of resistance mutations, a step that is not included in AMRPlusPlus 2.0 and it is available only for RGI and PointFinder, which however require assembled genomes or genes.

On semi-synthetic data, KARGVA has high accuracy in presence of non-resistance gene mutations, high error rates, and gene rearrangements. The multiple matching strategy increases ARGV detection accuracy, can report ARGVs that share high genetic similarity, and deal with configurations of resistance mutations not reported in literature. For instance, if a gene G develops resistance either through mutations {A, B, C} or {B, C, D}, and one sample presents with {A, C, D}, its *k*-mers align to G and pass the statistical assessment. However, one possible issue with this approach is that –even if sequences are collapsed– the ARGV database can still store the same ARGV twice or more if there are different laboratory-confirmed configurations of resistance mutations, e.g., using the example above, there would be two entries of for gene G considering both {A, B, C} and {B, C, D} mutations separately.

We also show that KARGVA presents very low false positive rates with respect to bacterial genes not necessarily involved in antimicrobial resistance, as well as specific mutant chromosomal genes that were found in antibiotic-susceptible samples. Although the semi-synthetic data are designed in a rigorous manner, the availability of standardized benchmark datasets from real experiments is auspicated, as discussed by Marini *et al*. in regards to ARG classifiers [27]

A limitation of the software is that the data structures, from a computational point of view, are not memory efficient. While the triple hash table design guarantees most search operations in constant time, there is considerable memory overhead in the padding of Java types/classes and legacy data structures (e.g., the String type and HashMap class). Also, the file parsing/writing is made with a standard BufferedReader and BufferedWriter, simply optimizing the buffer size, and the whole program is implemented serially. The KARGVA database is small, therefore the impact on processing times and memory usage is minimal, and KARGVA is about 6.5 times faster than AMRPlusPlus, independently from the FASTQ file size. Nonetheless, since new antibiotics and new ARGVs are discovered every year, it is advisable to foresee more efficient parsing and *k*-mer handling, considering also succinct data structures and parallelization [28]. Also, porting the software to mobile architectures –iOS or Android, and ARM chipsets– is warranted, given the growth of miniaturized, portable, point-of-care sequencing, like Nanopore MinION [29]. Since KARGVA is written in Java, the porting to mobile should not be a challenge (although consumer-grade applications have 512MB or 1GB memory limit depending on the operating system version), aside needs of optimization, and device overheating issues [30].

In the re-analysis of the MetaSUB data presented in Danko *et al*. [24], we confirm authors’ findings relative to percentages of unclassified reads (41% with KrakenUniq, in line with our results). We also confirm that –as ARG and ARGV databases grow– there is the expected increase in detection of antimicrobial resistance in the samples. The ARGV profiles retrieved by KARGVA seem more similar across cities than the ARG profiles by AMRPlusPlus. We find a strong correlation between number of ARGs and ARGVs found among samples; however, the median number of retrieved ARGVs is much larger than ARGs, even though ARG databases contain more gene entries. KARGVA substantially improves the ARGV finding rate with respect to other algorithms in the majority of the considered classes.

In accordance with Danko *et al*. [24], we find very low concordance between geographic distance among cities and distribution patterns of ARG/ARGVs, Further analysis adding the surface layer does not shed more insights, since there is high heterogeneity in the ARGV city-surface profile pairs. We cautionary do not want to draw any conclusion regarding ARGV patterns among cities, as we expect major unmeasured confounders, and we do not have a reference evolutionary history. The same in fact holds also if analyzing ARGs. To our knowledge, there are no established models to draw evolutionary relationships for metagenomes. Phylogenetic and phylodynamic trees at the species level could be inferred by assembling core genomes, possibly excluding any ARG or resistance mutation to remove bias from convergent evolution, although findings might not be insightful given that samples have been collected at the global level within a small time period –usually such analyses are meaningful for regional outbreaks [31]. Nonetheless, Danko *et al*. [24] were able to predict successfully geographic origin of samples using machine learners. Besides city-specific trends in taxa prevalence, ARG/ARGV patterns among cities might be associated to a plethora of factors, from population habits (diet, hygiene), ecological (cleaning schedules of public transportation system, characteristics of the users, e.g., youth, office workers, commuters from rural areas), public health practices (antibiotic usage guidelines and stewardship), in a mixture of common causes, mediators, and common effects for antibiotic resistance.

In conclusion, KARGVA provides reliable characterization of ARGVs, suitable for large metagenomics studies as well as targeted whole genome sequencing, and fills a current toolset and operational gap in a field where only a few limited options are available, with high potential for translational applications.

## Acknowledgment

This work was in part supported by US grants NIH NIAID R01AI145552, NIH NIAID R01AI141810, NSF SCH 2013998, and USDA AFRI 2019-67017-29110.

## Notes

https://github.com/DataIntellSystLab/KARGVA

### Competing Interest Statement

The authors have declared no competing interest.

## References

[1] C. J. Murray et al., “Global burden of bacterial antimicrobial resistance in 2019: a systematic analysis,” The Lancet, vol. 399, no. 10325, pp. 629–655, Feb. 2022, doi: 10.1016/S0140-6736(21)02724-0.

[2] C. D. Iwu, L. Korsten, and A. I. Okoh, “The incidence of antibiotic resistance within and beyond the agricultural ecosystem: A concern for public health,” MicrobiologyOpen, vol. 9, no. 9, p. e1035, Jul. 2020, doi: 10.1002/mbo3.1035.

[3] V. A. C. de Abreu, J. Perdigão, and S. Almeida, “Metagenomic Approaches to Analyze Antimicrobial Resistance: An Overview,” Front. Genet., vol. 0, 2021, doi: 10.3389/fgene.2020.575592.

[4] W. Gu, S. Miller, and C. Y. Chiu, “Clinical Metagenomic Next-Generation Sequencing for Pathogen Detection,” Annu. Rev. Pathol., vol. 14, pp. 319–338, Jan. 2019, doi: 10.1146/annurev-pathmechdis-012418-012751.

[5] M. Boolchandani, A. W. D’Souza, and G. Dantas, “Sequencing-based methods and resources to study antimicrobial resistance,” Nat. Rev. Genet., vol. 20, no. 6, pp. 356–370, Jun. 2019, doi: 10.1038/s41576-019-0108-4.

[6] B. P. Alcock et al., “CARD 2020: antibiotic resistome surveillance with the comprehensive antibiotic resistance database,” Nucleic Acids Res., vol. 48, no. D1, pp. D517–D525, Jan. 2020, doi: 10.1093/nar/gkz935.

[7] E. Doster et al., “MEGARes 2.0: a database for classification of antimicrobial drug, biocide and metal resistance determinants in metagenomic sequence data,” Nucleic Acids Res., vol. 48, no. D1, pp. D561–D569, Jan. 2020, doi: 10.1093/nar/gkz1010.

[8] “Database resources of the National Center for Biotechnology Information | Nucleic Acids Research | Oxford Academic.” https://academic.oup.com/nar/article/48/D1/D9/5585551 (accessed Oct. 26, 2021).

[9] V. Bortolaia et al., “ResFinder 4.0 for predictions of phenotypes from genotypes,” J. Antimicrob. Chemother., vol. 75, no. 12, pp. 3491–3500, Dec. 2020, doi: 10.1093/jac/dkaa345.

[10] X. Yin et al., “ARGs-OAP v2.0 with an expanded SARG database and Hidden Markov Models for enhancement characterization and quantification of antibiotic resistance genes in environmental metagenomes,” Bioinforma. Oxf. Engl., vol. 34, no. 13, pp. 2263–2270, Jul. 2018, doi: 10.1093/bioinformatics/bty053.

[11] M. Prosperi and S. Marini, “KARGA: Multi-platform Toolkit for k-mer-based Antibiotic Resistance Gene Analysis of High-throughput Sequencing Data,” in 2021 IEEE EMBS International Conference on Biomedical and Health Informatics (BHI), Jul. 2021, pp. 1–4. doi: 10.1109/BHI50953.2021.9508479.

[12] S. Marini et al., “AMR-meta: a k-mer and metafeature approach to classify antimicrobial resistance from high-throughput short-read metagenomics data,” GigaScience, vol. 11, p. giac029, Jan. 2022, doi: 10.1093/gigascience/giac029.

[13] S. M. Lakin et al., “Hierarchical Hidden Markov models enable accurate and diverse detection of antimicrobial resistance sequences,” Commun. Biol., vol. 2, Aug. 2019, doi: 10.1038/s42003-019-0545-9.

[14] G. Mk, F. Kj, and D. G, “Improved annotation of antibiotic resistance determinants reveals microbial resistomes cluster by ecology,” ISME J., vol. 9, no. 1, Jan. 2015, doi: 10.1038/ismej.2014.106.

[15] G. Arango-Argoty, E. Garner, A. Pruden, L. S. Heath, P. Vikesland, and L. Zhang, “DeepARG: a deep learning approach for predicting antibiotic resistance genes from metagenomic data,” Microbiome, vol. 6, Feb. 2018, doi: 10.1186/s40168-018-0401-z.

[16] B. Coculescu, “Antimicrobial resistance induced by genetic changes,” J. Med. Life, vol. 2, no. 2, pp. 114–123, Apr. 2009.

[17] I. Sultan, S. Rahman, A. T. Jan, M. T. Siddiqui, A. H. Mondal, and Q. M. R. Haq, “Antibiotics, Resistome and Resistance Mechanisms: A Bacterial Perspective,” Front. Microbiol., vol. 9, 2018, Accessed: Apr. 19, 2022. [Online]. Available: https://www.frontiersin.org/article/10.3389/fmicb.2018.02066

[18] M. Prosperi, M. Salemi, T. Azarian, F. Milicchio, J. A. Johnson, and M. Oliva, “Unexpected Predictors of Antibiotic Resistance in Housekeeping Genes of Staphylococcus Aureus,” ACM-BCB ACM Conf. Bioinforma. Comput. Biol. Biomed. ACM Conf. Bioinforma. Comput. Biol. Biomed., vol. 2019, pp. 259–268, Sep. 2019, doi: 10.1145/3307339.3342138.

[19] M. C. F. Prosperi, L. Prosperi, R. R. Gray, and M. Salemi, “On counting the frequency distribution of string motifs in molecular sequences,” Int. J. Biomath., vol. 05, no. 06, p. 1250055, May 2012, doi: 10.1142/S1793524512500556.

[20] N. A. O’Leary et al., “Reference sequence (RefSeq) database at NCBI: current status, taxonomic expansion, and functional annotation,” Nucleic Acids Res., vol. 44, no. D1, pp. D733–745, Jan. 2016, doi: 10.1093/nar/gkv1189.

[21] J. J. Davis et al., “The PATRIC Bioinformatics Resource Center: expanding data and analysis capabilities,” Nucleic Acids Res., vol. 48, no. D1, pp. D606–D612, Jan. 2020, doi: 10.1093/nar/gkz943.

[22] H. Gourlé, O. Karlsson-Lindsjö, J. Hayer, and E. Bongcam-Rudloff, “Simulating Illumina metagenomic data with InSilicoSeq,” Bioinformatics, vol. 35, no. 3, pp. 521–522, Feb. 2019, doi: 10.1093/bioinformatics/bty630.

[23] C. Mason et al., “The Metagenomics and Metadesign of the Subways and Urban Biomes (MetaSUB) International Consortium inaugural meeting report,” Microbiome, vol. 4, no. 1, p. 24, Jun. 2016, doi: 10.1186/s40168-016-0168-z.

[24] D. Danko et al., “A global metagenomic map of urban microbiomes and antimicrobial resistance,” Cell, vol. 184, no. 13, pp. 3376-3393.e17, Jun. 2021, doi: 10.1016/j.cell.2021.05.002.

[25] “Improved metagenomic analysis with Kraken 2 | Genome Biology | Full Text.” https://genomebiology.biomedcentral.com/articles/10.1186/s13059-019-1891-0 (accessed Dec. 04, 2021).

[26] S. Nurk, D. Meleshko, A. Korobeynikov, and P. A. Pevzner, “metaSPAdes: a new versatile metagenomic assembler,” Genome Res., vol. 27, no. 5, pp. 824–834, May 2017, doi: 10.1101/gr.213959.116.

[27] “Towards routine employment of computational tools for antimicrobial resistance determination via high-throughput sequencing | Briefings in Bioinformatics | Oxford Academic.” https://academic.oup.com/bib/article/23/2/bbac020/6536119?login=true (accessed Apr. 20, 2022).

[28] C. Marchet, C. Boucher, S. J. Puglisi, P. Medvedev, M. Salson, and R. Chikhi, “Data structures based on k-mers for querying large collections of sequencing data sets,” Genome Res., Dec. 2020, doi: 10.1101/gr.260604.119.

[29] M. Oliva, F. Milicchio, K. King, G. Benson, C. Boucher, and M. Prosperi, “Portable nanopore analytics: are we there yet?,” Bioinforma. Oxf. Engl., vol. 36, no. 16, pp. 4399–4405, Aug. 2020, doi: 10.1093/bioinformatics/btaa237.

[30] F. Milicchio and M. Prosperi, “Experimental Survey on Power Dissipation of k-mer-Handling Data Structures for Mobile Bioinformatics,” in 2021 IEEE International Conference on Bioinformatics and Biomedicine (BIBM), Dec. 2021, pp. 3201–3206. doi: 10.1109/BIBM52615.2021.9669768.

[31] M. Prosperi et al., “Molecular Epidemiology of Community-Associated Methicillin-resistant Staphylococcus aureus in the genomic era: a Cross-Sectional Study,” Sci. Rep., vol. 3, p. 1902, May 2013, doi: 10.1038/srep01902.

