## Supplementary Tables 1-2 for "The *K*-mer Antibiotic Resistance Gene Variant Analyzer (KARGVA)"

### V. Supplementary material

*Supplementary Table 1:*

*MetaSUB Output term correspondence in KARGVA (MEGARes class used a reference)*

| CARD/RGI | MEGARes class | NDARO | Notes |
|---|---|---|---|
| - | Oxazolidinone | OXAZOLIDINONE | |
| aminocoumarin | Aminocoumarins | AMINOCOUMARIN | |
| aminoglycosides | Aminoglycosides | AMINOGLYCOSIDE | |
| amoxicillin | betalactams | BETA-LACTAM | |
| basR | Cationic_antimicrobial_peptides | COLISTIN | Megares group: BASRS; merge with Lipopetides |
| beta-lactam | betalactams | BETA-LACTAM | |
| colistin | Lipopeptides | COLISTIN | Megares group: BASRS; merge with Lipopetides |
| D-Ala-D-Ala ligase | Glycopeptides | - | |
| dapsone | Sulfonamides | SULFONAMIDE | |
| daptomycin | Lipopeptides | LIPOPEPTIDE | |
| eatAv | Multi-drug_resistance | - | Megares group: EATAV |
| elfamycin | Elfamycins | - | |
| ethambutol | Mycobacterium_tuberculosis-specific_Drug | - | Megares mechanism: Ethambutol_resistant_arabinosyltransferase |
| ethionamide | Mycobacterium_tuberculosis-specific_Drug | - | Megares mechanism: Ethionamide-resistant_mutant |
| fluoroquinolones | Fluoroquinolones | QUINOLONES | |
| fosfomycin | Fosfomycin | FOSFOMYCIN | |
| Fusidic_acid | Fusidic_acid | FUSIDIC_ACID | |
| GE2270A | Elfamycins | - | |
| imipenem | betalactams | BETA-LACTAM | |
| isoniazid | Mycobacterium_tuberculosis-specific_Drug | ISONIAZID/TRICLOSAN | Megares mechanism: Isoniazid-resistant_mutant |
| kirromycin | Elfamycins | - | |
| lysocin | Lipopeptides | LIPOPEPTIDE | |
| Moxifloxacin | Fluoroquinolones | QUINOLONES | |
| mtrR | Drug_and_biocide_MFS_efflux_regulator | - | |
| mupirocin | Mupirocin | - | |
| nitrofurantoin | - | NITROFURAN | |
| Omp36 | betalactams | BETA-LACTAM | |

| | | | |
|---|---|---|---|
| para-aminosalicylic acid | Mycobacterium_tuberculosis-specific_Drug | - | Megares mechanism: Para-aminosalicylic_acid_resistant_mutant |
| PhoP | Lipopeptides | COLISTIN | Merged with Cationic_antimicrobial_peptides |
| PIB (por) | Multi-drug_resistance | - | Not specific at the class level, could be "MDR_mutant_porin_proteins" |
| pleuromutilin | Pleuromutilin | - | |
| prothionamide | - | - | |
| prothionamide | - | - | |
| Pulvomycin | Elfamycins | - | |
| pyrazinamide | Pyrazinamide-resistant_mutant | - | |
| ramR | Multi-drug_RND_efflux_regulator | EFFLUX | Megares group: RAMR |
| rifampicin | Rifampin | RIFAMYCIN | |
| rpld | MLS | MACROLIDE | |
| rpsJ | Tetracyclines | TETRACYCLINE | |
| soxR | Drug_and_biocide_resistance | EFFLUX | Megares group: SOXR |
| soxS | Drug_and_biocide_and_metal_resistance | - | Megares group: SOXS |
| streptomycin | Aminoglycosides | AMINOGLYCOSIDE | |
| sulfonamides | Sulfonamides | SULFONAMIDE | |
| triclosan | Phenolic_compound_resistance | TRICLOSAN | |
| triclosan | Phenolic_compound_resistance | QUINOLONE/TRICLOSAN | |
| vancomycin | Glycopeptides | - | |
| zoliflodacin | - | - | |

*Supplementary Table 2:*

*MetaSUB output term correspondence between PointFinder and MEGARes class*

| PointFilder | MEGARes class | Notes |
|---|---|---|
| AMIKACIN | Aminoglycosides | |
| Amoxicillin | betalactams | |
| Ampicillin | betalactams | |
| Azithromycin | MLS | |
| BEDAQUILINE | - | Not considered for class-specific comparison (not present in MEGARes) |
| CAPREOMYCIN | Aminoglycosides | |
| carbapenem | betalactams | |
| cefixime | betalactams | |
| cefotaxime | betalactams | |
| cefoxitin | betalactams | |
| ceftazidime | betalactams | |
| cephalosporins | betalactams | |
| Ciprofloxacin | Fluoroquinolones | |
| Clindamycin | MLS | |
| CLOFAZIMINE | Fluoroquinolones | |
| Colistin | Lipopeptides | |
| CYCLOSERINE | - | Not considered for class-specific comparison (not present in MEGARes) |
| Erythromycin | MLS | |
| ETHAMBUTOL | Mycobacterium_tuberculosis-specific_Drug | Not considered for class-specific comparison (described in MEGARes as Mechanism) |
| ETHIONAMIDE | Mycobacterium_tuberculosis-specific_Drug | Not considered for class-specific comparison (described in MEGARes as Mechanism) |
| fluoroquinolone | Fluoroquinolones | |
| FLUOROQUINOLONE | Fluoroquinolones | |
| Gentamicin C | Aminoglycosides | |
| ISONIAZID | Mycobacterium_tuberculosis-specific_Drug | Not considered for class-specific comparison (described in MEGARes as Mechanism) |
| KANAMYCIN | Aminoglycosides | |

| | | |
|---|---|---|
| Kanamycin A | Aminoglycosides | |
| Kasugamycin | Aminoglycosides | |
| Linezolid | Oxazolidinone | |
| Nalidixic acid | Fluoroquinolones | |
| Neomycin | Aminoglycosides | |
| norfloxacin | Fluoroquinolones | |
| Para-aminosalicyclic acid | Mycobacterium_tuberculosis-specific_Drug | Not considered for class-specific comparison (described in MEGARes as Mechanism) |
| Paromomycin | Aminoglycosides | |
| piperacillin | betalactams | |
| PYRAZINAMIDE | Mycobacterium_tuberculosis-specific_Drug | Not considered for class-specific comparison (described in MEGARes as Mechanism) |
| Rifampicin | Rifampin | |
| RIFAMPICIN | Rifampin | |
| Spectinomycin | Aminoglycosides | |
| Streptomycin | Aminoglycosides | |
| STREPTOMYCIN | Aminoglycosides | |
| Sulfamethoxazole | Sulfonamides | |
| Telithromycin | MLS | |
| Tetracycline | Tetracyclines | |
| tigecycline | Tetracyclines | |
| Tobramycin | Aminoglycosides | |
| XDR-TB | - | Not considered for class-specific comparison (not present in MEGARes) |
| Penicillins | betalactams | |
| Macrolides | MLS | |
| Quinolones | Fluoroquinolones | |
| Sulphadoxine | Sulfonamides | |
| Pyrimethamine | - | Not considered for class-specific comparison (not present in MEGARes) |
| Chloroquine | - | Not considered for class-specific comparison (not present in MEGARes) |
| Amodiaquine | - | Not considered for class-specific comparison |

| | | |
|---|---|---|
| | | (not present in MEGARes) |
| Piparaquine | - | Not considered for class-specific comparison (not present in MEGARes) |
| Lumefantrine | - | Not considered for class-specific comparison (not present in MEGARes) |
| Mefloquine | - | Not considered for class-specific comparison (not present in MEGARes) |
| Artemisinin | - | Not considered for class-specific comparison (not present in MEGARes) |
| fusidic acid | Fusidic_acid | |
| Trimethoprim | Trimethoprim | Not considered for class-specific comparison (not present in KARGVA) |
| Mupirocin | Mupirocin | |
| Cefazolin | betalactams | |
| Ceftriaxone | betalactams | |
| Ceftaroline | betalactams | |
